# Variantkey: A Reversible Numerical Representation of Human Genetic Variants

**DOI:** 10.1101/473744

**Authors:** Nicola Asuni, Steven Wilder

## Abstract

Human genetic variants are usually represented by four values with variable length: chromosome, position, reference and alternate alleles. There is no guarantee that these components are represented in a consistent way across different data sources, and processing variant-based data can be inefficient because four different comparison operations are needed for each variant, three of which are string comparisons. Existing variant identifiers do not typically represent every possible variant we may be interested in, nor they are directly reversible. Similarly, genomic regions are typically represented inconsistently by three or four values. Working with strings, in contrast to numbers, poses extra challenges on computer memory allocation and data-representation. To overcome these limitations, a novel reversible numerical encoding schema for human genetic variants (*VariantKey*) and genomics regions (*RegionKey*), is presented here alongside a multi-language open-source software implementation (https://github.com/Genomicsplc/variantkey). *VariantKey* and *RegionKey* represents variants and regions as single 64 bit numeric entities, while preserving the ability to be searched and sorted by chromosome and position. The individual components of short variants can be directly read back from the *VariantKey*, while long variants are supported with a fast lookup table.

## 1 Introduction

There is no standard guideline for consistent representation of genetic variants. For a given genome assembly, different classes of genetic variants are typically represented in the *Variant Call Format* (VCF) [1] as a combination of four components: chromosome name, base position, reference allele and alternate allele(s). The alternate field can contain comma-separated strings for multiallelic variants but they can be decomposed into biallelic ones.

Since there is no guarantee that the variant components are represented in a consistent way across different data sources, a normalization step is also required. Even with normalized variants, basic operations can be inefficient due to the necessity to perform four different comparison operations for each variant, three of which are string comparisons. Working with strings poses extra challenges on storage, memory allocation and data-representation due to their variable-length nature.

Existing variant identifiers don’t provide a solution to this general problem because they are typically assigned, they require an expensive lookup operation to be retrieved or reversed and they don’t represent every possible variant we may be interested in. In particular, the *dbSNP* reference identifier (rs#, “rs ID” or “rs tag”) [2] can’t uniquely identify a variant in a given reference genome.

To overcome these limitations, VariantKey, a novel reversible numerical encoding schema for human genetic variants, is presented here. VariantKey represents variants as a single 64 bit numeric entity, while preserving the ability to be searched and sorted by chromosome and position. The individual components of short variants can be directly read back from the VariantKey, while long variants are supported with a fast lookup table.

Similarly, regions of the human genome are often written as three or four components: chromosome, start-position, end-position and sometimes strand. The meaning of the start and end position differs between applications. RegionKey is a novel reversible numerical encoding schema for human genomic regions, where the components of the region can be directly read from the RegionKey.

A reference implementation of VariantKey, RegionKey and utility functions is provided as a multi-language software library (https://github.com/Genomicsplc/variantkey) released under the open source MIT license. For portability and performance, the core source code is written in header-only C programming language in a way that is also compatible with `C++`. Full wrappers for Go, Python, R and a basic implementation in Javascript are also provided. The source code is documented and fully covered by unit tests.

## 2 Human Genetic Variant Definition

A genetic variant is a difference from the reference DNA nucleotide sequence.

In this context, the human genetic variant for a given genome assembly is defined as the set of four components compatible with the VCF format:

- **CHROM** - *chromosome*: An identifier from the reference genome. It only has 26 valid values: autosomes from 1 to 22, the sex chromosomes X=23 and Y=24, mitochondria MT=25 and a symbol NA=0 to indicate an invalid value.
- **POS** - *position*: The reference position in the chromosome, with the first base having position 0. The largest expected value is less than 250 million [3] to represent the last base pair in Chromosome 1.
- **REF** - *reference base(s)*: String containing a sequence of reference nucleotide letters. The value in the POS field refers to the position of the first base in the String.
- **ALT** - *alternate base(s)*: Single alternate non-reference allele. String containing a sequence of nucleotide letters. Multiallelic variants must be decomposed in individual biallelic variants.

## 3 Variant Decomposition and Normalization

### 3.1 Decomposition

In the Variant Call Format (VCF) the alternate field can contain comma-separated strings for multiallelic variants, while in this context we only consider biallelic variants to allow for allelic comparisons between different data sets.

For example, the multiallelic variant:

`{CHROM=1, POS=3759889, REF=TA, ALT=TAA,TAAA,T}`

can be decomposed as three biallelic variants:

`{CHROM=1, POS=3759889, REF=TA, ALT=TAA}`
`{CHROM=1, POS=3759889, REF=TA, ALT=TAAA}`
`{CHROM=1, POS=3759889, REF=TA, ALT=T}`

In VCF files the decomposition from multiallelic to biallelic variants can be performed using the `vt` software tool [4] with the command:

`vt decompose -s source.vcf -o decomposed.vcf`

The “-s” option (smart decomposition) splits up INFO and GENOTYPE fields that have number counts of R and A appropriately [5].

### 3.2 Normalization

A normalization step is required to ensure a consistent and unambiguous representation of variants.

As shown in Figure 1, there are multiple ways to represent the same variant, but only one can be considered “normalized” as defined by [6]:

- A variant representation is normalized if and only if it is left aligned and parsimonious.
- A variant representation is left aligned if and only if its base position is smallest among all potential representations having the same allele length and representing the same variant.
- A variant representation is parsimonious if and only if the entry has the shortest allele length among all VCF entries representing the same variant.

**Figure 1:**
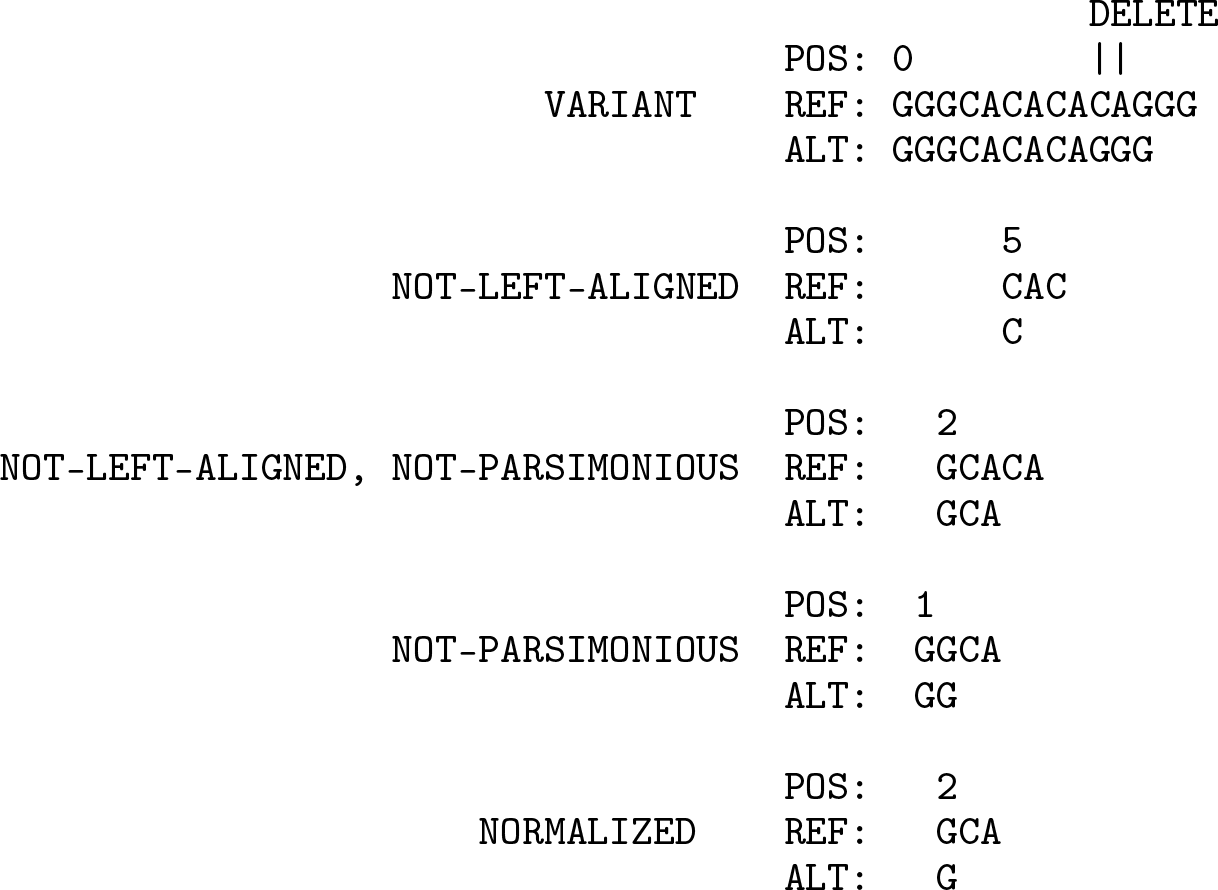
Example of entries representing the same variant.

In VCF files the variant normalization can be performed using the vt software tool [7]:

`vt normalize -m -r genome.fa -o normalized.vcf decomposed.vcf`

or the bcftools command [8]:

`bcftools norm -f genome.fa -o normalized.vcf decomposed.vcf`

or decompose and normalize with a single command:

`bcftools norm --multiallelics -any -f genome.fa -o normalized.vcf source.vcf`

#### 3.2.1 Normalization Function

Individual biallelic variants can be normalized using the `normalize_variant` function provided by the *variantkey* library. The `normalize_variant` function first checks if the reference allele matches the genome reference. The match is considered valid and consistent if there is a perfect letter-by-letter match, and valid but not consistent if one or more letter matches an equivalent one. Equivalent IUPAC nucleotide codes for one or more base pairs are accepted, as defined in Table 1.

**Table 1:** Single-letter nucleotide codes [9]

| SYMBOL | DESCRIPTION | EQ. BASES | COMPLEMENT |
| --- | --- | --- | --- |
| A | Adenine | A | T |
| C | Cytosine | C | G |
| G | Guanine | G | C |
| T | Thymine | T | A |
| W | Weak | A T | W |
| S | Strong | C G | S |
| M | aMino | A C | K |
| K | Keto | G T | M |
| R | puRine | A G | Y |
| Y | pYrimidine | C T | R |
| B | not A (B comes after A) | C G T | V |
| D | not C (D comes after C) | A G T | H |
| H | not G (H comes after G) | A C T | D |
| V | not T (V comes after T and U) | A C G | B |
| N | aNy base (not a gap) | A C G T | N |

If the reference allele is not valid, the `normalize_variant` function tries to find a reference match with one of the following variant transformations:

- swap the reference and alternate alleles - *sometimes it is not clear which one is the reference and which one is the alternate allele*.
- flip the alleles letters (use the complement letters) - *sometimes the alleles refers to the other DNA strand*.
- swap and flip.

Note that the *swap* and *flip* processes can lead to false positive cases, especially when considering *Single Nucleotide Polymorphisms* (SNP). The return code of the `normalize_variant` function can be used to discriminate or discard variants that are not consistent.

If the variant doesn’t match the genome reference, then the original variant is returned with an error code.

If both alleles have length 1, the normalization is complete and the variant is returned. Otherwise, a custom implementation of the vt normalization algorithm [6] is applied:

- while break, do
  – if any of the alelles is empty and the position is greater than zero, then
    ∗ extend both alleles one letter to the left using the nucleotide in the corresponding genome reference position;
  – else
    ∗ if both alleles end with the same letter and they have length 2 or more, then
      · truncate the rightmost letter of each allele;
    ∗ else
      · break (exit the while loop);
- while both alleles start with the same letter and have length 2 or more, do
  – truncate leftmost letter of each allele;

The genome reference binary file (`fasta.bin`), used by the `normalize_variant` function, can be obtained from a FASTA file using the `resources/tools/fastabin.sh` script. This script extracts only the first 25 sequences for chromosomes 1 to 22, X, Y and MT. Precompiled versions can be downloaded from: https://sourceforge.net/projects/variantkey/files/.

#### 3.2.2 Normalized VariantKey

The variantkey library provides the normalized_variantkey function that returns the VariantKey of the normalized variant. This function should be used instead of the variantkey function if the input variant is not normalized.

## 4 VariantKey Format

For a given reference genome the VariantKey format encodes a Human Genetic Variant (CHROM, POS, REF and ALT) as 64 bit unsigned integer number (8 bytes or 16 hexadecimal symbols).

If the variant has not more than 11 bases between REF and ALT, the correspondent VariantKey can be directly reversed to get back the individual CHROM, POS, REF and ALT components.

If the variant has more than 11 bases, or non-base nucleotide letters are contained in REF or ALT, the VariantKey can be fully reversed with the support of a binary lookup table.

The VariantKey format doesn’t represent universal codes, it only encodes CHROM, POS, REF and ALT, so each code is unique for a given reference genome. The direct comparisons of two VariantKeys makes sense only if they both refer to the same genome reference.

The VariantKey is composed of 3 sections arranged in 64 bit:

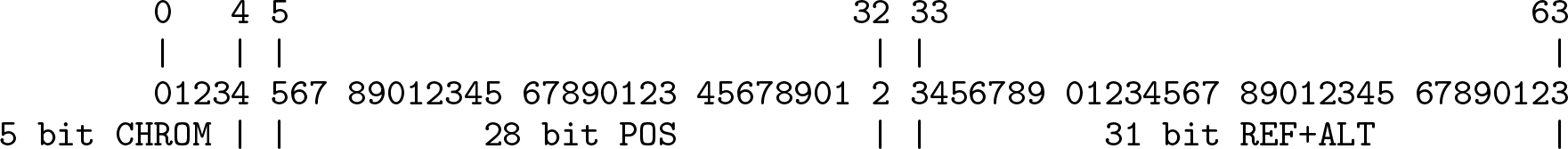

**Table 2:** Example of VariantKey encoding

|  | CHROM | POS | REF | ALT |
| --- | --- | --- | --- | --- |
| **Raw variant** | chr19 | 29238770 | TC | TG |
| **Normalized variant** | 19 | 29238771 | C | G |
| **VariantKey bin** | 10011 | 0001101111100010010111110011 | 0001 0001 01 10 00000000000000000000 |  |
| **VariantKey hex** | 98DF12F988B00000 |  |  |  |
| **VariantKey dec** | 11015544076520914944 |  |  |  |

### 4.1 CHROM Encoding

The chromosome is encoded as unsigned integer number: 1 to 22, X=23, Y=24, MT=25, NA=0. This section is 5 bit long, so it can store up to 2^5^ = 32 symbols, enough to contain the required 25 canonical chromosome symbols + NA. The largest value is: 25 dec = 19 hex = 11001 bin. Values from 26 to 31 are currently reserved. They can be used to indicate 6 alternative modes to interpret the remaining 59 bit. For instance, one of these values can be used to indicate the encoding of variants that occurs in non-canonical contigs.

**Figure 2:**
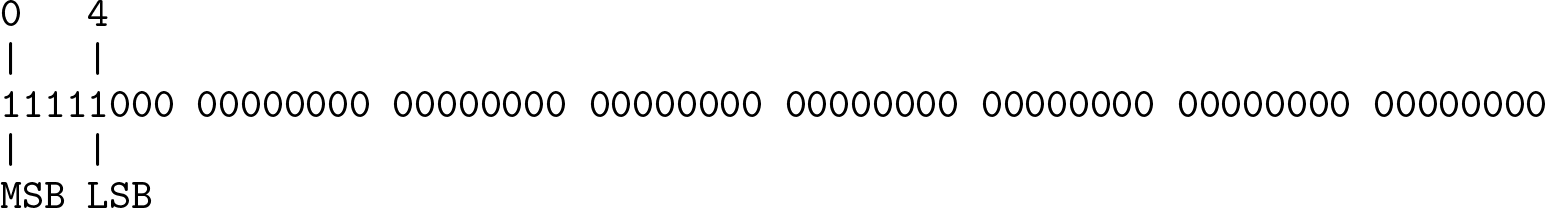
CHROM binary mask (F800000000000000 hex = 17870283321406128128 dec)

Example: `‘chr19’ str = 19 dec = 10011 bin`

### 4.2 POS Encoding

The reference position in the chromosome is encoded as unsigned integer number with the first base having position 0. This section is 28 bit long, so it can store up to 2^28^ = 268, 435, 456 symbols, enough to contain the maximum position found in the largest human chromosome.

**Figure 3:**
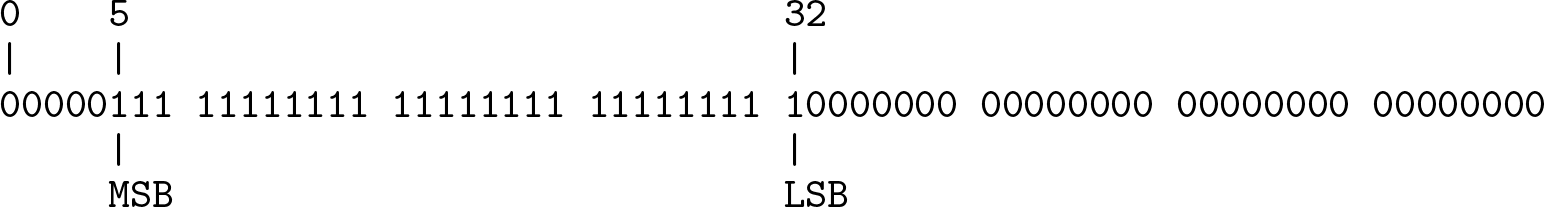
POS binary mask (7FFFFFF80000000 hex = 576460750155939840 dec)

Example: `29238771 dec = 0001101111100010010111110011 bin`

### 4.3 REF+ALT Encoding

The reference and alternate alleles are encoded in 31 bit.

This section allow two different type of encodings:

**Figure 4:**
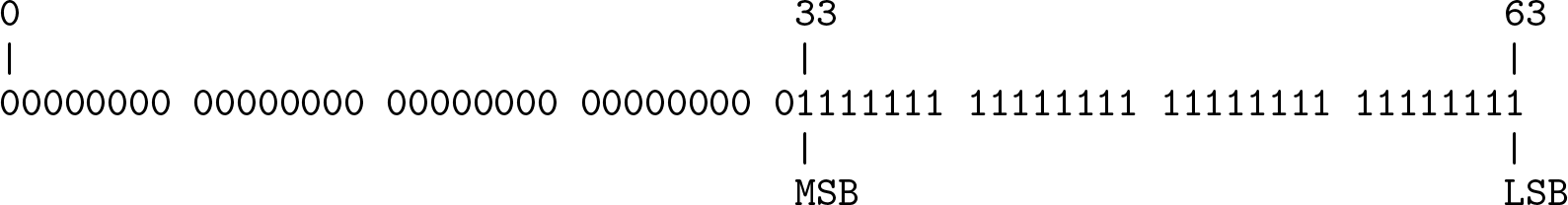
REF+ALT binary mask (7FFFFFFF hex = 2147483647 dec)

#### 4.3.1 Non-reversible encoding

If the total number of nucleotides between REF and ALT is more then 11, or if any of the alleles contains nucleotide letters other than base A, C, G and T, then the LSB (least significant bit) is set to 1 and the remaining 30 bit are filled with an hash value of the REF and ALT strings.

The hash value is calulated using a custom fast non-cryptographic algorithm based on *MurmurHash3* [10]. A lookup table (`nrvk.bin`) is required to reverse the REF and ALT values.

In the normalized dbSNP VCF file GRCh37.p13.b150 there are only 0.365% (1,229,769 / 337,162,128) variants that requires this encoding. Amongst those, the maximum number of variants that share the same chromosome and position is 15. With 30 bit the probability of hash collision is approximately 10^−7^ for 15 elements, 10^−6^ for 46 and 10^−5^ for 146. The size of the non-reversible lookup table for dbSNP GRCh37.p13.b150 is only 45.7MB, so it can be easily contained in memory.

#### 4.3.2 Reversible encoding

If the total number of nucleotides between REF and ALT is 11 or less, and they only contain base letters A, C, G and T, then the LSB is set to 0 and the remaining 30 bit are used as follows:

- bit 1-4 indicate the number of bases in REF; the capacity of this section is 2^4^ = 16 but the maximum expected value is 10 dec = 1010 bin;
- bit 5-8 indicate the number of bases in ALT; the capacity of this section is 2^4^ = 16 but the maximum expected value is 10 dec = 1010 bin;
- the following 11 groups of 2 bit are used to represent REF bases followed by ALT, with the following encoding:

– `A = 0 dec = 00 bin`
– `C = 1 dec = 01 bin`
– `G = 2 dec = 10 bin`
– `T = 4 dec = 11 bin`

Examples:

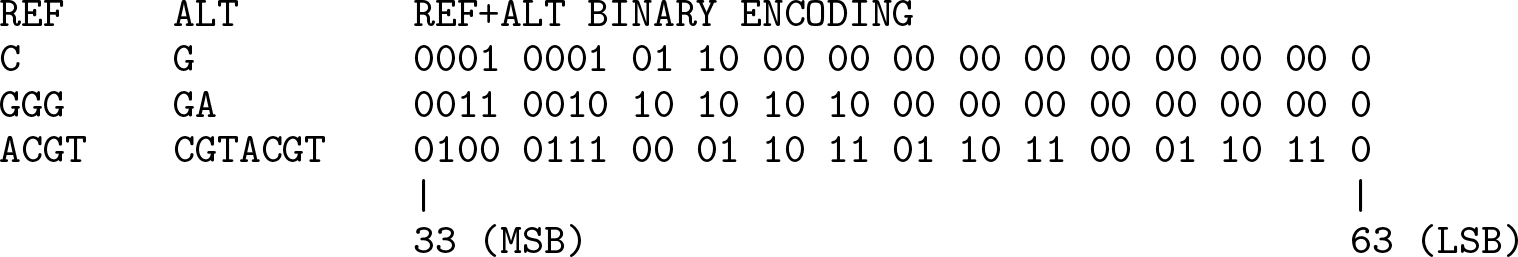

The reversible encoding covers 99.635% of the variants in the normalized dbSNP VCF file GRCh37.p13.b150.

## 5 RegionKey Format

For a given reference genome the RegionKey format encodes a Human Genomic Region (CHROM, STARTPOS, ENDPOS, STRAND) as a 64 bit unsigned integer number (8 bytes or 16 hexadecimal symbols). A RegionKey can be directly reversed to get back the individual CHROM, STARTPOS, ENDPOS and STRAND components.

The RegionKey format doesn’t represent universal codes, it only encodes CHROM, STARTPOS, ENDPOS and STRAND, so each code is unique for a given reference genome. The direct comparisons of two RegionKeys makes sense only if they both refer to the same genome reference.

The RegionKey is composed of 4 sections arranged in 64 bit:

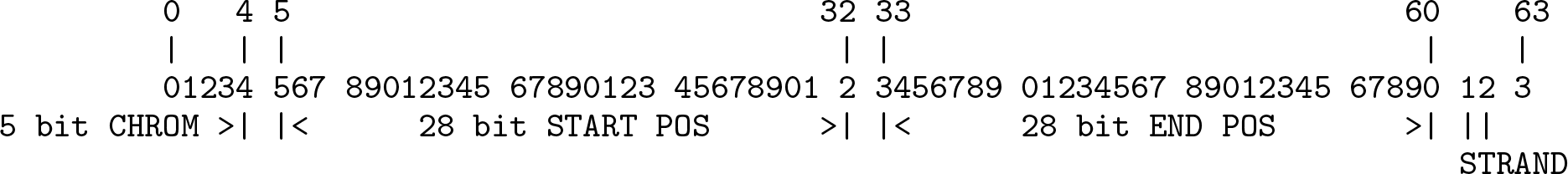

**Table 3:**
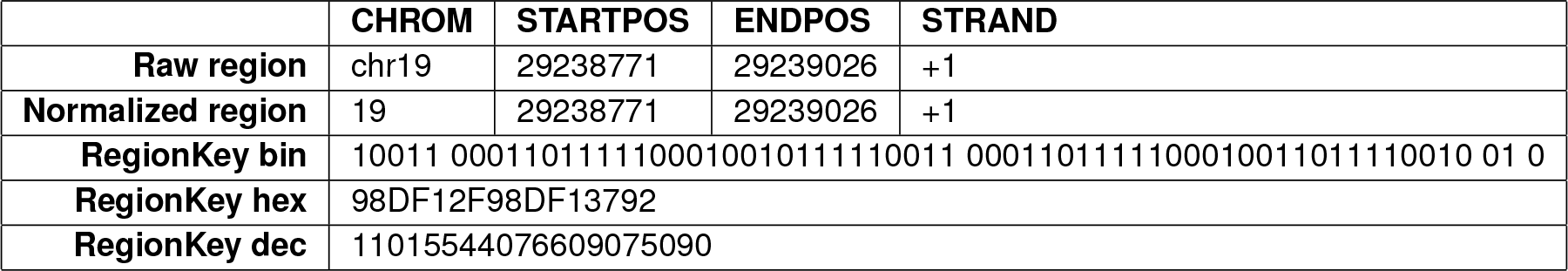
Example of RegionKey encoding

### 5.1 CHROM Encoding

The encoding for chromosome is the same as for VariantKey, see 4.1.

### 5.2 STARTPOS Encoding

The encoding for the region start position is the same as for POS in VariantKey, see 4.2.

### 5.3 ENDPOS Encoding

The region end position in the chromosome is encoded as unsigned integer number with the first reference base having position 0. This section is 28 bit long, so it can store up to 2^28^ = 268, 435, 456 symbols, enough to contain the maximum position found in the largest human chromosome. The end position is equivalent to (STARTPOS + REGION_LENGTH), such that the base having position ENDPOS is not included in the region.

**Figure 5:**
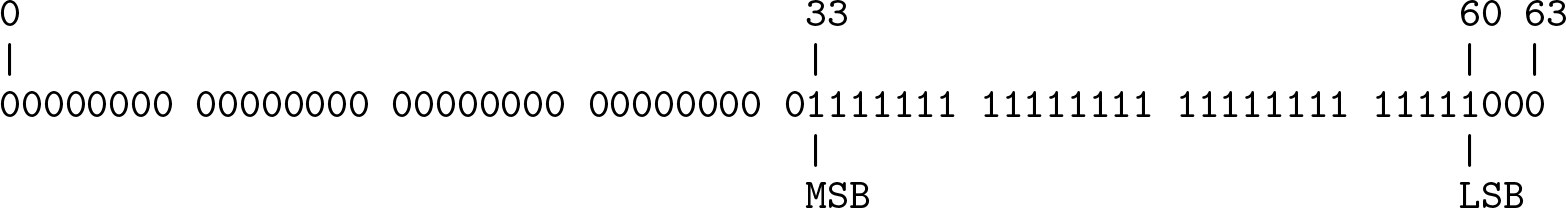
ENDPOS binary mask (7FFFFFF8 hex = 2147483640 dec)

Example: `29239026 dec = 0001101111100010011011110010 bin`

### 5.4 STRAND Encoding

Encodes the DNA strand direction. This is useful when encoding genic regions.

**Figure 6:**
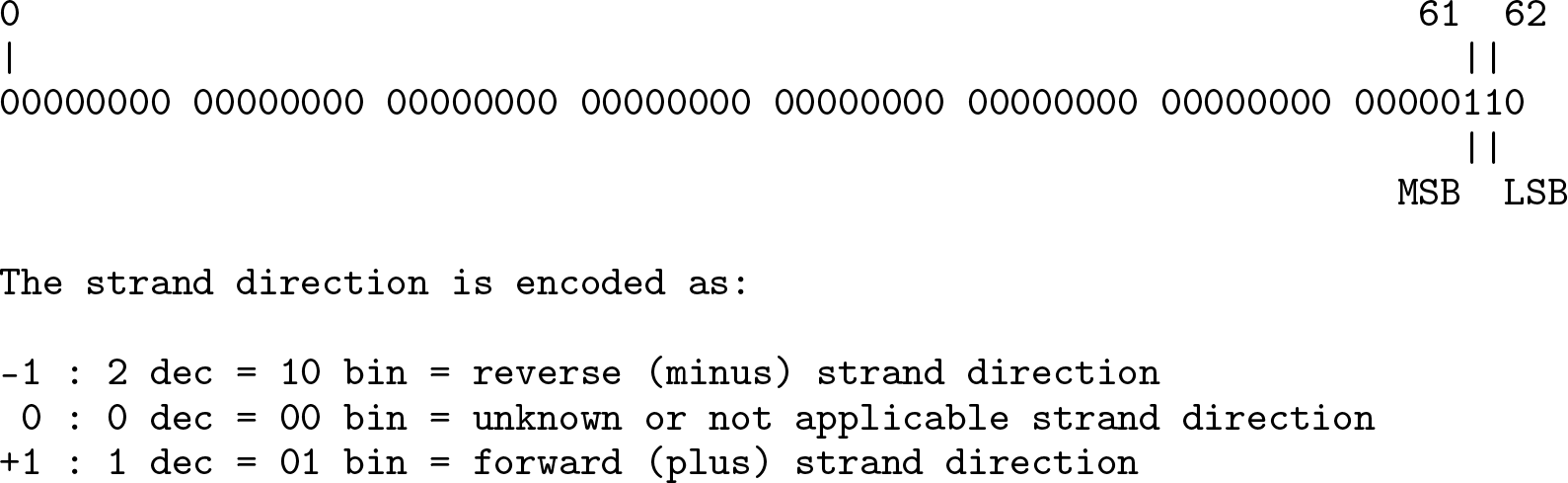
STRAND binary mask (3 hex = 3 dec)

## 6 VariantKey and RegionKey Properties

- They can be encoded and decoded on-the-fly.
- Sorting by VariantKey (or RegionKey) is equivalent of sorting by CHROM and POS (CHROM and START-POS).
- The 64 bit VariantKey or RegionKey can be exported as a single 16 character hexadecimal string.
- Sorting the hexadecimal representation of VariantKey or RegionKey in alphabetical order is equivalent of sorting numerically.
- Comparing two variants by VariantKey only requires comparing two 64 bit numbers, a very well optimized operation in current computer architectures. In contrast, comparing two normalized variants in VCF format requires comparing one numbers and three strings.
- VariantKey or RegionKey can be used as a main database key to index data by “variant” or “region”. This simplify common searching, merging and filtering operations.
- All types of database joins between two data sets (inner, left, right and full) can be easily performed using the VariantKey or RegionKey as index.
- When CHROM, REF and ALT are the only strings in a table, replacing them with VariantKey allows to work with numeric only tables with obvious advantages. This also allows to represent the data in a compact binary format where each column uses a fixed number of bit or bytes, with the ability to perform a quick binary search on the first sorted column.

## 7 Example VariantKey Application

A direct application of the VariantKey representation is the ability to be used in binary lookup table files.

The variantkey library provides tools to create and use binary lookup table files to convert rsID to VariantKey and vice-versa:

- `rsvk.bin`
  Lookup table to retrieve VariantKey from rsID.
  This binary file can be generated by the `resources/tools/rsvk.sh` script from a TSV file.
  This can also be in *Apache Arrow* file format with a single *RecordBatch*, or *Feather* format. The first column must contain the rsID sorted in ascending order.
- `vkrs.bin`
  Lookup table to retrieve rsID from VariantKey.
  This binary file can be generated by the `resources/tools/vkrs.sh` script from a TSV file.
  This can also be in *Apache Arrow* file format with a single *RecordBatch*, or *Feather* format. The first column must contain the VariantKey sorted in ascending order.

Software benchmarks on the variantkey library, performed using a laptop with 16GB of RAM and Intel core i7 7th gen CPU, shows that it is possible to retrieve from a file with 400M items, a VariantKey from rsID or viceversa in about 48ns, or more than 20M items per second.

## 8 Overlapping Regions

Two regions `A` and `B` overlap if the following condition is satisfied:

- `(A_CHROM = B_CHROM) and (A_STARTPOS < B_ENDPOS) and (A_ENDPOS > B_STARTPOS)`

Given a list `L` of sorted RegionKeys, and the maximum region length `L_MAX_REGION_LENGTH` of the entries in `L`, the following algorithm can be used to efficiently find which entries overlap a defined region `R`:

- Binary search only on the CHROM and STARTPOS bits:
  – find the list upper bound (`UB`), the maximal entry in `L`, such that:
    ∗ `(L_CHROM = R_CHROM) and (L_STARTPOS < R_ENDPOS)`
  – find the list lower bound (`LB`), the minimal entry less than `UB`, such that:
    ∗ `(L_CHROM = R_CHROM) and (L_STARTPOS > (R_STARTPOS - L_MAX_REGION_LENGTH))`
- Linear search between `LB` and `UB` entries for (`L_ENDPOS > R_STARTPOS`).

**Figure 7:**
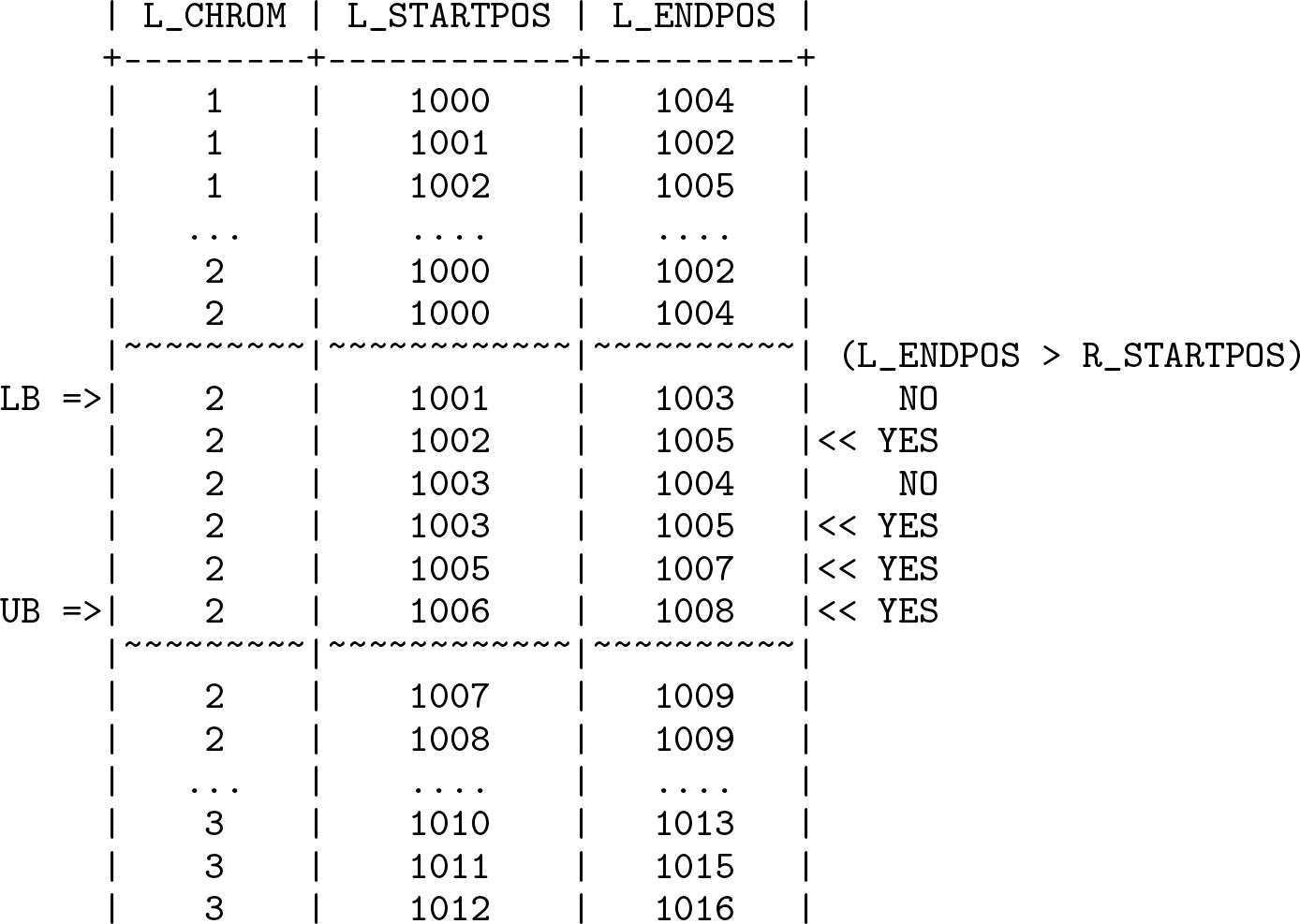
Overlapping regions (`R_CHROM=2, R_STARTPOS=1004, R_ENDPOS=1007, L_MAX_REGION_LENGTH=4`)

## 9 Conclusions

Existing human genetic variant or genomic region representation formats are not designed to maximize computational performance and do not provide a general solution to efficiently represent any arbitrary variant. VariantKey, a new reversible numerical encoding schema for human genetic variants, and RegionKey for human genomic regions are proposed here, with a common publicly available implementation. These new computer-friendly formats are expected to improve the performance of large-scale computation of genetic data.

